# Tracing the Evolutionary Histories of Human Cholinergic Proteins Across Eukaryotes: A Comparative Phylogenomic Study

**DOI:** 10.64898/2026.11.04.764462

**Authors:** Ankit Thakur, Mahesh Kulharia

## Abstract

Acetylcholine signalling depends on a coordinated network of enzymes, transporters, and receptors whose evolutionary histories need not be congruent. Here, we examined 26 curated human cholinergic reference proteins across 89 selected eukaryotic proteomes using a human-reference-anchored phylogenomic framework. Candidate sequences were identified through sequence similarity searches and evaluated using domain architecture, orthology assessment, and family-level phylogenetic analyses. Gene-tree/species-tree reconciliation was performed for selected datasets to investigate putative duplication and loss histories. The results revealed heterogeneous patterns among the investigated protein groups. CHAT- and SLC18A3-associated candidates were recovered across multiple sampled bilaterian lineages, whereas SLC5-family-related sequences showed broader representation, including sampled ctenophores, cnidarians, and bilaterians. Within the latter group, a distinct vertebrate-associated subgroup was identified in relation to human SLC5A7. Cholinesterase analyses recovered an echinoderm-associated grouping and distinct vertebrate ACHE- and BCHE-associated groups, although their deeper evolutionary relationship received comparatively lower support. Muscarinic and nicotinic receptor analyses recovered selected subtype- and subunit-associated groupings, but limited taxonomic representation and sequence instability constrained broader evolutionary interpretation. Collectively, these findings reveal heterogeneous taxonomic distributions and phylogenetic relationships among proteins corresponding to the human cholinergic system. The study provides a comparative framework for examining these protein groups while emphasizing that their recovered evolutionary histories must be interpreted in the context of reference-sequence selection, taxonomic sampling, and phylogenetic resolution.

## Introduction

Acetylcholine is a signalling molecule with central roles in neuronal communication and physiological regulation. In vertebrates, it mediates transmission at neuromuscular junctions and autonomic ganglia and is released by parasympathetic neurons at many peripheral targets. In the central nervous system, cholinergic projections modulate neuronal activity across distributed circuits, contributing to cognitive processes that depend on coordinated signalling (Haga, 2013; Ananth *et al*., 2023). These diverse functions depend on a molecular system that controls transmitter availability and couples its release to distinct cellular responses. Understanding the evolutionary relationships among the components of this system can therefore illuminate how conserved biochemical processes and diversified receptors contribute to cholinergic signalling.

Cholinergic neurotransmission relies on coordinated synthesis, transport, receptor activation and hydrolysis. Choline O-acetyltransferase (ChAT; encoded by *CHAT*) synthesizes acetylcholine, whereas the high-affinity choline transporter CHT1 (SLC5A7) supports choline uptake for transmitter production and recycling. The vesicular acetylcholine transporter VAChT (SLC18A3) loads acetylcholine into secretory vesicles, making it available for regulated release (Erickson *et al*., 1994; Ennis and Blakely, 2016). After release, acetylcholine activates nicotinic and muscarinic receptors and is hydrolysed principally by acetylcholinesterase (AChE); the related butyrylcholinesterase (BChE) can also hydrolyse acetylcholine and contribute to its clearance in particular contexts (Mesulam *et al*., 2002; Pezzementi, Nachon and Chatonnet, 2011). Synthesis, choline uptake, vesicular loading and degradation are thus related but molecularly distinguishable processes.

The two principal acetylcholine receptor classes couple the same ligand to responses through different protein architectures. Nicotinic acetylcholine receptors belong to the Cys-loop ligand-gated ion-channel superfamily and assemble as pentamers, with each subunit contributing four transmembrane helices. Their neuronal and muscle-associated subunits form receptor combinations with differing signalling properties; the structure of the human α4β2 receptor illustrates the organization of a heteromeric neuronal receptor (Morales-Perez, Noviello and Hibbs, 2016; Jiao *et al*., 2019). Muscarinic receptors, by contrast, are class A G-protein-coupled receptors with seven transmembrane helices. The five mammalian subtypes, M1–M5, differ in their preferred coupling: M1, M3 and M5 generally signal through Gq/11, whereas M2 and M4 preferentially engage Gi/o proteins (Haga, 2013; Pedersen, Bergqvist and Larhammar, 2018). Their contrasting structures and modes of signalling make these receptor families complementary subjects for evolutionary comparison.

Comparative studies indicate that cholinergic signalling has a history across animal lineages, but they also argue against treating its molecular components as a single evolutionary unit. In the cnidarian *Nematostella vectensis*, receptor expression and pharmacological experiments support neuronal and non-neuronal roles for nicotinic acetylcholine receptors, while phylogenetic analysis indicates diversification in both cnidarian and bilaterian lineages (Faltine-Gonzalez and Layden, 2019). Among bilaterians, comparative genomics has revealed pronounced, lineage-specific expansion of nicotinic receptor genes in molluscs (Jiao *et al*., 2019). Vertebrate muscarinic receptors show a different pattern: sequence and genomic-context analyses support diversification of their subfamilies in association with early vertebrate genome duplications (Pedersen, Bergqvist and Larhammar, 2018). These observations make it important to examine enzymes, transporters, ion channels and G-protein-coupled receptors individually, rather than assume that their present-day association reflects a shared history of origin or diversification.

Existing comparative work provides foundations for such an investigation, although its emphases and taxonomic scopes differ. Studies of the amphioxus cholinergic locus have characterized *ChAT* and *VAChT* homologues and their expression, while broader solute-carrier analyses have examined transporter-family evolution across eukaryotes (Candiani *et al*., 2008; Höglund *et al*., 2011). Cholinesterase evolution has also been investigated through comparative characterization of vertebrate enzymes, including an atypical BChE from medaka (Pezzementi, Nachon and Chatonnet, 2011). Together with receptor-focused analyses, these studies illuminate particular components, but their different gene sets and sampling frameworks leave the relationships among the human cholinergic components less readily comparable across a shared taxonomic context. Moreover, duplication and subsequent divergence complicate the distinction between orthologues and paralogues, so a functional annotation or sequence match alone does not establish evolutionary correspondence (Koonin, 2005).

A human-reference-anchored analysis offers a consistent starting point for addressing this comparability problem. Independently using curated human protein sequences as queries allows candidate homologues to be considered in relation to defined components of the human system, while retaining the possibility that their evolutionary histories differ. Sequence similarity can be evaluated alongside domain architecture and family-specific phylogenies to distinguish plausible homologous relationships from misleading annotations or assignments within diversified families. Where gene trees are sufficiently resolved, comparison with a species tree can further inform interpretations of speciation, duplication and loss (Page and Charleston, 1997). Such an approach traces lineages associated with human references; it does not by itself reconstruct the complete ancestral cholinergic system or establish the absolute origin of each family. Similarly, failure to recover a candidate from a sampled proteome is not, on its own, proof of evolutionary absence.

Here, we investigate the distribution and phylogenetic relationships of 26 curated human cholinergic reference proteins across 89 selected eukaryotic proteomes using a human-reference-anchored comparative phylogenomic framework. The reference set encompasses proteins associated with acetylcholine biosynthesis, catabolism, transport, and receptor-mediated signalling. Our objective is to characterize the taxonomic representation of candidate homologues, examine their relationships within the corresponding protein families, and explore patterns of diversification using gene-tree/species-tree reconciliation where the available phylogenetic datasets permit. By applying a common analytical framework to these distinct molecular components, we aim to compare their recovered evolutionary distributions and highlight differences in their phylogenetic resolution and diversification patterns. Importantly, this approach traces sequence lineages associated with selected human reference proteins within the sampled proteomes and does not seek to reconstruct the complete ancestral cholinergic repertoire or establish the absolute evolutionary origin of each component.

## Materials and Methods

This study explores the evolutionary emergence and distribution of selected human cholinergic reference genes using computational approaches to examine proteins associated with acetylcholine biosynthesis, transport, catabolism and regulation, as well as receptor-mediated functions across diverse organisms. All protein sequence and proteomic data were retrieved from publicly accessible databases, including NCBI and UniProtKB. The UniProt FASTA sequences used as query proteins and the proteome FASTA files employed for local BLAST database construction are provided in Online Resource A. To facilitate local BLAST database construction and avoid potential issues arising from complex FASTA headers, sequence headers were masked, and corresponding ID-mapping dictionaries were generated to enable the retrieval of original sequence descriptions and annotations (Online Resource B). As this study relied exclusively on publicly available protein sequence and proteome datasets and did not involve human participants or experimental animals, ethical approval was not required.

### Species Selection

Proteome datasets for the selected species were primarily retrieved from NCBI and UniProtKB, supplemented by additional publicly accessible repositories. Initially, 107 proteomes representing major metazoan lineages and selected non-metazoan eukaryotes were compiled to provide a broader phylogenetic context. This sampling strategy was designed to ensure broad taxonomic coverage for identifying homologous sequences corresponding to the 26 human reference proteins, rather than to comprehensively reconstruct gene content or evolutionary histories across all eukaryotic lineages.

Proteome completeness was evaluated using BUSCO v6.1.1 across all 107 proteomes, employing the Eukaryota lineage dataset (eukaryota_odb10), which comprises 255 single-copy orthologs (Simão *et al*., 2015). BUSCO was applied as a comparative quality-control measure rather than an absolute indicator of genome or proteome completeness. Proteomes satisfying the predefined completeness threshold established in the original analysis were retained, yielding a final dataset of 89 proteomes with BUSCO completeness scores of ≥80%. However, BUSCO scores may vary depending on lineage-specific gene content and the representation of conserved single-copy orthologs; consequently, lower scores do not necessarily reflect overall incompleteness of the underlying genome or proteome. Accordingly, the BUSCO threshold was used exclusively as a quality-control criterion and not as definitive evidence of genome completeness. The distribution of retained proteomes across taxonomic groups, along with their corresponding BUSCO completeness scores, is presented in Supplementary Table 1.

### Protein selection and homology search

Human reference protein sequences used for homology searches were obtained from UniProtKB/Swiss-Prot (The UniProt Consortium, 2023) and supplemented with curated information from the relevant literature. Each human protein was treated as an independent sequence anchor for its respective homology search. The 26 focal human reference proteins constituted the primary seed queries used to search across the 89 proteomes. Homology searches were performed using PSI-BLAST v2.12.0+ (Altschul *et al*., 1997), with each of the 26 curated human reference proteins submitted as an independent query. Each search was conducted for three iterations using an E-value threshold of 1 × 10□^2^□, and the resulting candidate sequences were subsequently subjected to downstream filtering procedures (Online Resource C). The homology-search strategy was designed to identify candidate sequences related to the selected human reference proteins rather than to comprehensively recover all members of the corresponding protein families. Given the use of human proteins as individual sequence anchors and the stringent search threshold applied, recovery was expected to favour candidates exhibiting detectable sequence similarity to the human references under the specified search conditions. Consequently, failure to recover a candidate was interpreted as non-detection within the present analytical framework rather than definitive evidence of evolutionary absence.

Filtered candidate sequences were functionally annotated using BLASTP (McGinnis and Madden, 2004) against the UniProtKB/Swiss-Prot database (The UniProt Consortium, 2023) (Online Resource D). The top-scoring hit was considered suitable for putative functional annotation when sequence coverage exceeded 80% and percentage identity was greater than 40% (Rost, 1999; Pearson, 2013). These functional annotations were used as supporting evidence and were not considered a replacement for sequence homology assessment, domain architecture analysis, or phylogenetic placement (Rost, 1999).

Redundant protein sequences within individual species were removed using CD-HIT v4.8.1 (Huang *et al*., 2010), applying a sequence identity threshold of 100% (-c 1.0). This procedure removed identical protein entries while retaining sequence variation among candidate homologues. Broccoli v1.3 was subsequently employed to organize candidate sequences into putative orthologous groups using phylogenetic and similarity-based information (Derelle, Philippe and Colbourne, 2020). These assignments were considered supporting evidence for the evolutionary classification of candidate sequences and were interpreted alongside the corresponding family-level phylogenies and domain annotations. Accordingly, sequences associated with individual human reference proteins are referred to as human-reference-associated homologues unless their specific orthologous relationship is supported by the broader phylogenetic context. The corresponding Broccoli assignments are provided in Online Resource F.

Candidate cholinergic receptor sequences were assessed for predicted transmembrane helices using Phobius v1.01 (Käll, Krogh and Sonnhammer, 2007). Considering the expected architecture of nicotinic acetylcholine receptor subunits, which generally contain four transmembrane helices, and muscarinic acetylcholine receptors, which contain seven, sequences with 3-8 predicted transmembrane helices were retained for downstream analyses. This criterion was used as an initial structural plausibility filter rather than as definitive evidence of receptor identity or orthology. Domain annotations and functional classifications were additionally examined using InterProScan (Blum *et al*., 2025), with the corresponding results provided in Online Resource G. Receptor sequence redundancy was subsequently reduced using CD-HIT with an identity threshold of 80% (-c 0.8).

Sequence-space relationships among candidate proteins were explored using CLANS2 v2.2.2 (Frickey and Lupas, 2004). Candidate sequences from biosynthetic, catabolic, receptor, and transporter categories were analysed independently using pairwise similarity-based clustering. Clustering was examined across P-value thresholds ranging from 1 × 10 ¹ to 1 × 10 ¹ to visualize relationships among candidate sequences and assist in organizing family-specific datasets (Online Resource H). CLANS2 clustering was treated as an exploratory approach for examining sequence relationships, rather than as independent evidence of orthology or evolutionary lineage assignment. The resulting datasets were subsequently examined using multiple sequence alignment and phylogenetic reconstruction.

### Phylogenetic Tree Construction Analysis

Following homology searches, domain-architecture assessment, and CLANS2-based filtering, phylogenetic datasets were constructed at the protein-family or orthogroup level, depending on the evolutionary relationships under investigation. To examine relationships among human paralogues, the corresponding human reference proteins and their identified homologues were analysed together within family-level phylogenies. This strategy was applied to the catabolic enzyme family comprising ACHE and BCHE, as well as the muscarinic and nicotinic receptor subunit families. Each phylogenetic dataset was independently aligned using MAFFT v7.505 (Katoh, Rozewicki and Yamada, 2019). Following alignment, poorly aligned and potentially phylogenetically uninformative regions were removed using trimAl v1.5.rev0 (Capella-Gutiérrez, Silla-Martínez and Gabaldón, 2009), with positions containing more than 70% gaps excluded (Online Resource I).

Maximum-likelihood phylogenies were inferred using IQ-TREE2 v2.2.2.7 (Minh *et al*., 2020). ModelFinder (Kalyaanamoorthy *et al*., 2017) was used to identify the best-fitting amino acid substitution model according to the Bayesian Information Criterion. Branch support was evaluated using 1,000 ultrafast bootstrap replicates (UFBoot) and Transfer Bootstrap Expectation

(TBE) values based on 100 non-parametric bootstrap replicates (Hoang *et al*., 2018; Zaharias, Lemoine and Gascuel, 2023). These two support measures were interpreted independently as complementary assessments of phylogenetic support. In particular, differences between UFBoot and TBE values were considered when evaluating the stability and interpretation of individual clades. High branch-support values were not treated as independent evidence of gene orthology or functional conservation. Maximum-likelihood gene trees were inferred as unrooted phylogenies and subsequently supplied to GeneRax for reconciliation with the rooted species tree.

The UFB analysis incorporated the specified substitution-model and rate-heterogeneity search settings, together with branch-length reporting, nearest-neighbour interchange optimization, and additional branch-support assessments. The corresponding IQ-TREE analysis parameters were: -m MFP, -mset cpREV,Dayhoff,DCMut,FLU,HIVb,HIVw,JTT,JTTDCMut,LG,mtART,mtMAM,mtREV,mtZO A,mtMet,mtVer,mtInv,Poisson,PMB,rtREV,VT,WAG,GTR20, -mrate E,I,G,I+G,R, -B 1000, --wbtl, --bnni, --alrt 1000, and --abayes. TBE support was estimated using -b 100 --tbe. The resulting IQ-TREE phylogenies are provided in Online Resource J. Maximum-likelihood gene trees were inferred as unrooted phylogenies; consequently, no specific outgroup was designated during tree construction. For subsequent gene-tree/species-tree reconciliation, these unrooted maximum-likelihood gene trees were supplied to GeneRax (Morel *et al*., 2020) together with an independently constructed rooted species tree, as described below.

### Rogue Taxa Analysis

To identify phylogenetically unstable sequences that could potentially influence tree reconstruction and downstream evolutionary inferences, we applied two complementary approaches: the t-index (Lemoine *et al*., 2018) and the Leaf Stability Index (LSI) (Wilkinson, 2006). Implemented in IQ-TREE2, the t-index measures the frequency with which a taxon changes its phylogenetic position across bootstrap replicates, with higher values indicating greater topological instability. TBE-based t-index scores were primarily used in this study, and sequences with t-index values greater than 2 were classified as unstable. The Leaf Stability Index (LSI) provides an additional measure of taxon stability by evaluating how consistently a sequence retains its phylogenetic placement across bootstrap trees. LSI values were calculated using RogueNaRok v1.0.1 with TBE-based trees (Aberer, Krompass and Stamatakis, 2013), and sequences with low LSI score were considered unstable. Taxa identified as unstable by either the t-index or LSI analysis were excluded from the gene trees subsequently used for reconciliation to minimize their potential influence on downstream evolutionary interpretations. Importantly, phylogenetic instability was treated as an indication of uncertainty in sequence placement rather than evidence that a candidate sequence was necessarily incorrect or biologically absent. The complete identifiers of sequences excluded from reconciliation are provided in Online Resource K.

### Reconciliation Analysis

Gene-tree/species-tree reconciliation was performed using GeneRax v2.1.3 under the UndatedDL model (-r UndatedDL) to investigate potential gene duplication and loss scenarios within the sampled protein families (Morel *et al*., 2020). Reconciliation was performed independently for datasets retaining a sufficient number of phylogenetically usable sequences following the stability assessment. Datasets in which all terminal sequences were classified as unstable were not subjected to reconciliation.

For each dataset, the corresponding maximum-likelihood gene tree and a rooted species tree derived from the Open Tree of Life (OToL) were supplied to GeneRax (Hinchliff *et al*., 2015). The OToL topology was pruned to include the taxa represented in the corresponding phylogenetic dataset. The rooted species tree used for reconciliation is provided in Online Resource L.

The UndatedDL model evaluates gene-tree/species-tree compatibility through duplication and loss events. Consequently, the present reconciliation analyses were restricted to these evolutionary event classes and were not designed to evaluate horizontal gene transfer. The inferred duplication and loss events were interpreted as model-dependent scenarios conditional on the input gene trees, species-tree topology, and taxonomic representation. In particular, inferred loss events were not considered definitive evidence of gene absence in the corresponding evolutionary lineages.

Reconciled trees were visualized using ThirdKind v3.13.5 (Penel *et al*., 2022). The reconciliation visualizations and corresponding GeneRax outputs are provided in Online Resources M(a) and M(b), respectively. Ancestral placements were interpreted only within the context of the sampled species tree and were not considered estimates of absolute evolutionary origins or temporally calibrated divergence events.

Throughout this study, sequence counts refer to retained protein sequence entries or phylogenetic terminal tips and should not necessarily be interpreted as equivalent to the number of distinct genomic loci or independently validated gene copies. Similarly, duplication and loss events reported from gene-tree/species-tree reconciliation represent model-inferred evolutionary events under the specified analytical framework.

## Results

### Biosynthesis

#### Evolution of the Cholinergic Biosynthetic Machinery

Using human choline O-acetyltransferase (ChAT; UniProt: P28329) as the reference query, candidate homologous sequences passing the applied search and filtering criteria were recovered from 24 of the 89 sampled proteomes. These sequences represented selected bilaterian lineages, including vertebrates, echinoderms, arthropods, rotifers, and spiralians. No candidate sequence passed the applied identification and filtering criteria in the sampled non-bilaterian or non-metazoan proteomes. Phylogenetic reconstruction recovered a strongly supported vertebrate-associated grouping (TBE = 0.99, UFB = 97), an echinoderm-associated grouping (TBE = 1.00, UFB = 100), and a spider/tick cluster (TBE = 0.94, UFB = 89). In contrast, the grouping of rotifer and spiralian sequences received comparatively lower support (TBE = 0.67, UFB = 40), indicating uncertainty in their deeper phylogenetic relationship (Fig. 1A; Online Resource J). The stability assessment identified two potentially unstable sequences originating from *Centruroides vittatus* and *Danio rerio* (Online Resource K).

**Fig. 1.**
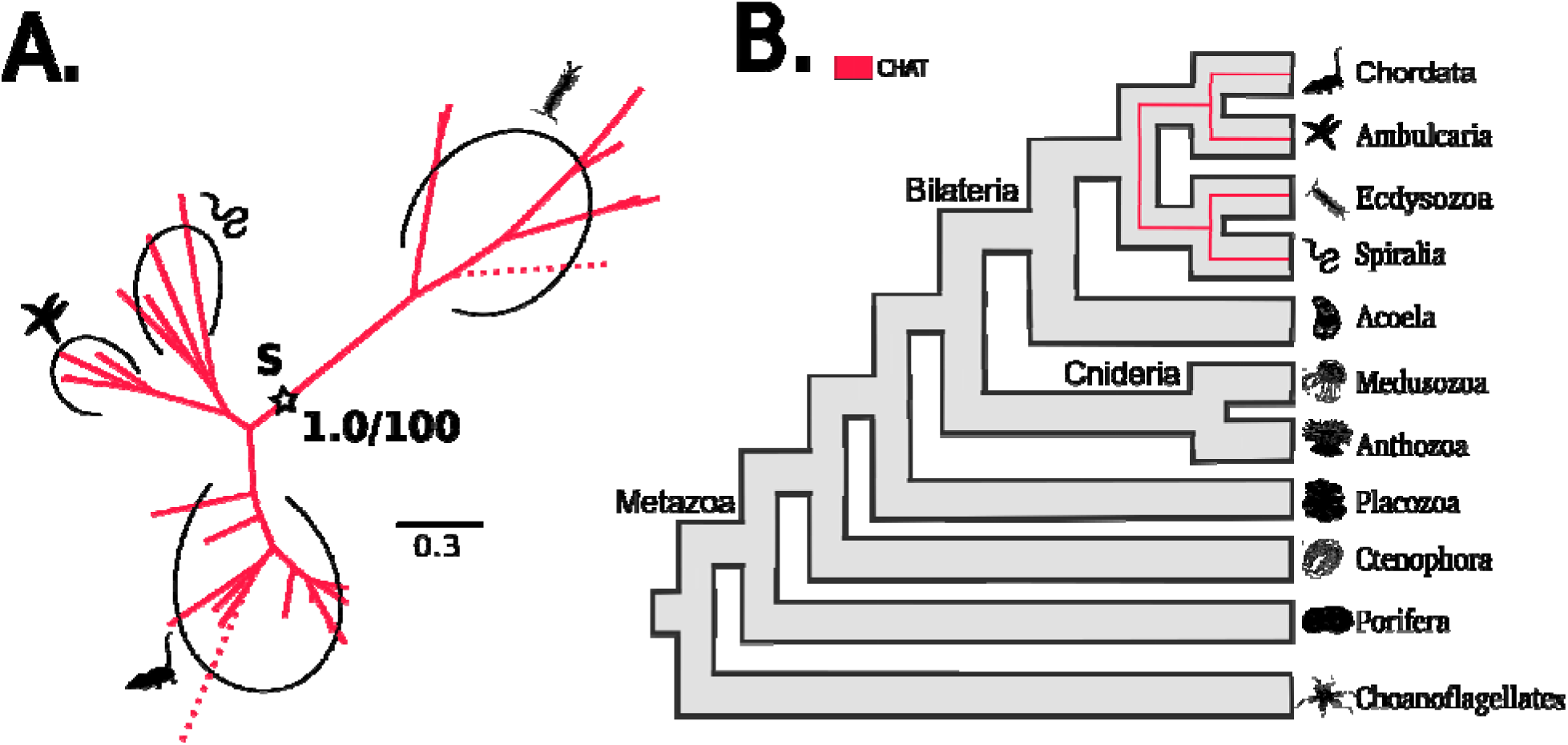
Phylogenetic relationships and gene-tree-species-tree reconciliation of choline O-acetyltransferase (CHAT). (A) Transfer bootstrap expectation tree showing the evolutionary relationships of the CHAT protein family. Nodal support values are shown as transfer bootstrap expectation (TBE; decimal values) and ultrafast bootstrap (UFB; whole-number percentages) for selected nodes. Dashed branches indicate sequences identified as unstable by the t-index or leaf stability index (LSI) analyses. Asterisks (✰) indicate speciation (S) or duplication (D) events, with TBE and UFB support values provided where applicable (B) Simplified representations of the corresponding gene-tree-species-tree reconciliations inferred using GeneRax. Species silhouettes were obtained from PhyloPic (https://www.phylopic.org/). Complete annotated phylogenetic trees and nodal support values are provided in Online Resource J, and detailed GeneRax reconciliation outputs are provided in Online Resource M(b).

Following the exclusion of two potentially unstable sequences, the CHAT reconciliation dataset comprised 33 terminal sequence tips. GeneRax reconciliation under the UndatedDL model identified nine duplication events and three loss events within the sampled dataset (Fig. 1B; Online Resources M(a) and M(b)). The inferred duplication events were mapped to branches associated with *Mus musculus, Gallus gallus, Anolis carolinensis, Takifugu rubripes, Patiria miniata, Centruroides vittatus,* and *Dermacentor albipictus*, whereas loss events were mapped to lineages associated with *Latimeria chalumnae, Lottia gigantea*, and *Capitella teleta*. These events represent reconciliation-based evolutionary scenarios under the supplied gene-tree/species-tree framework and should not be interpreted as independently confirmed gene-copy changes or definitive evidence of gene loss. The present analysis did not resolve an ancestral duplication separating distinct human CHAT-associated paralogous lineages.

### TRANSPORTERS

#### Evolution of the Vesicular Acetylcholine Transporter

Using human VAChT (UniProt: Q16572; encoded by SLC18A3, transports cytosolic acetylcholine into synaptic vesicles, enabling its storage and subsequent release during cholinergic neurotransmission) as the reference query, homologous sequences associated with the SLC18 transporter orthogroup were identified in 19 of the 89 sampled proteomes. Their distribution spanned diverse bilaterian lineages, including vertebrates, echinoderms, arthropods, molluscs and annelids. No homologues were recorded in the sampled non-bilaterian or non-metazoan proteomes among the sampled proteomes. Phylogenetic reconstruction recovered a vertebrate-associated clade (TBE = 0.91, UFB = 79), within which the mammalian sequences formed a strongly supported group (TBE = 1.00, UFB = 100), including a well-supported human-mouse grouping (TBE = 1.00, UFB = 100) (see Fig. 2A(red); Online Resource J).

**Fig. 2.**
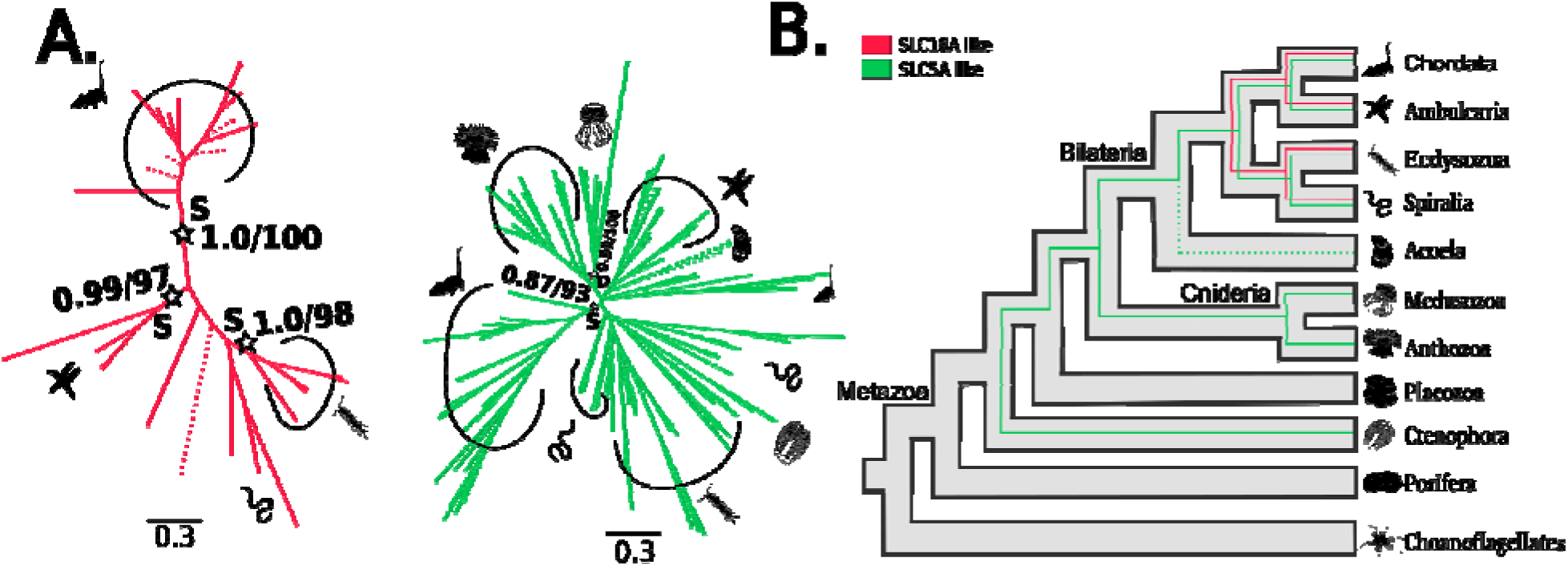
Phylogenetic relationships and gene-tree-species-tree reconciliation of Vesicular Acetylcholine Transporter (SLC18A3 / VAChT) and High-Affinity Choline Transporter 1 (SLC5A7 / CHT1). (A) Transfer bootstrap expectation (TBE) phylogenetic trees depicting the evolutionary relationships of the SLC18A and SLC5A protein families. Nodal support values are shown as transfer bootstrap expectation (TBE; decimal values) and ultrafast bootstrap (UFB; whole-number percentages) for selected nodes. Dashed branches indicate sequences identified as unstable by the t-index or leaf stability inde (LSI) analyses. Asterisks (✰) indicate speciation (S) or duplication (D) events, with TBE and UFB support values provided where applicable. (B) Simplified, combined representations of the corresponding gene-tree-species-tree reconciliations inferred using GeneRax. Species silhouettes were obtained from PhyloPic (https://www.phylopic.org/). Complete annotated phylogenetic trees and nodal support values are provided in Online Resource J, and detailed GeneRax reconciliation outputs are provided in Online Resource M(b).

The stability assessment identified seven potentially unstable sequences originating from *Xenopus tropicalis, Lethenterium reissneri, Latimeria chalumnae, Anolis carolinensis, Petromyzon marinus, Adineta vaga*, and *Callorhinchus milii*. These sequences were excluded from subsequent reconciliation, reducing the dataset from 30 initial phylogenetic tips to 23 retained tips. GeneRax reconciliation identified three duplication events and two loss events within the retained SLC18-associated dataset (Fig. 2B; Online Resources M(a) and M(b)). The inferred events included a duplication assigned to a node within the sampled bilaterian portion of the species tree, together with additional duplications in *Mus musculus* and *Takifugu rubripes*. Loss events were mapped to branches associated with Centruroides vittatus and the lineage leading to *Gallus gallus* and *Anolis carolinensis*. These findings provide a model-dependent representation of SLC18-associated sequence diversification among the sampled bilaterians rather than a definitive reconstruction of the complete evolutionary history of the transporter family.

#### Evolution of the High-Affinity Choline Transporter 1

Using human high-affinity choline transporter 1 (CHT1; UniProt: Q9GZV3), encoded by SLC5A7, as the reference query, candidate sequences associated with the broader SLC5 transporter family were recovered from 46 of the 89 sampled proteomes. These sequence showed a broader taxonomic representation than the SLC18A3-associated candidates, including sampled cnidarians, ctenophores, echinoderms, arthropods, molluscs, annelids, and chordates.

Phylogenetic reconstruction recovered a strongly supported vertebrate-associated subgroup comprising 18 sequences from 12 vertebrate species (TBE = 1.00, UFB = 100). Within this subgroup, the human, mouse, and opossum sequences formed a supported mammalian grouping (TBE = 0.94, UFB = 89), while the human–mouse grouping received TBE = 0.87 and UFB = 87. These relationships support the association of the vertebrate subgroup with the human SLC5A7 reference sequence. However, the broader phylogenetic backbone remained uncertain, with one 107-sequence grouping receiving TBE = 0.73 and UFB = 3, indicating substantial discordance between the two branch-support assessments (Fig. 2A(green); Online Resource J).

The stability assessment identified four potentially unstable sequence tips originating from Symsagittifera roscoffensis (two sequences), Strongylocentrotus purpuratus, and Corallochytrium limacisporum. Their exclusion reduced the reconciliation dataset from 196 initial phylogenetic tips to 192 retained tips. The C. limacisporum sequence was therefore not considered reliable evidence for the presence of this transporter-associated lineage in the sampled non-metazoan taxon.

GeneRax reconciliation under the UndatedDL model identified 120 duplication events and 51 loss events across the broader SLC5-associated dataset (Fig. 2B; Online Resources M(a) and M(b)). The reported lineage-associated expansions included Takifugu rubripes (13 duplication events), Anneissia japonica (12), Dermacentor albipictus (10), Lingula unguis (9), and Montipora foliosa (8). These counts describe model-inferred events across the analysed SLC5-associated sequence dataset and should not be interpreted as confirmed SLC5A7-specific duplication events. Taken together, the results indicate broad taxonomic representation of SLC5-family-related sequences, alongside a more clearly resolved vertebrate subgroup associated with human CHT1.

### Catabolism

#### Evolution of the Cholinesterases

Using human acetylcholinesterase (ACHE; UniProt: P22303; responsible for terminating cholinergic neurotransmission through rapid hydrolysis of acetylcholine (ACh) into choline and acetate) and butyrylcholinesterase (BCHE; UniProt: P06276; a related cholinesterase with broader substrate specificity that also hydrolyses ACh and other choline esters) as reference queries, we investigated the evolutionary relationships of the cholinesterase family across the 89 sampled proteomes. Homologues were detected in 27 species, spanning both major bilaterian lineages, including vertebrates, echinoderms, arthropods, cephalochordates, molluscs and annelids. No homologues were detected in the sampled non-bilaterian or non-metazoan proteomes. The phylogenetic analyses recovered a distinct echinoderm-associated clade (TBE = 0.93, UFB = 89) (see Fig. 3A; Online Resource J).

**Fig. 3.**
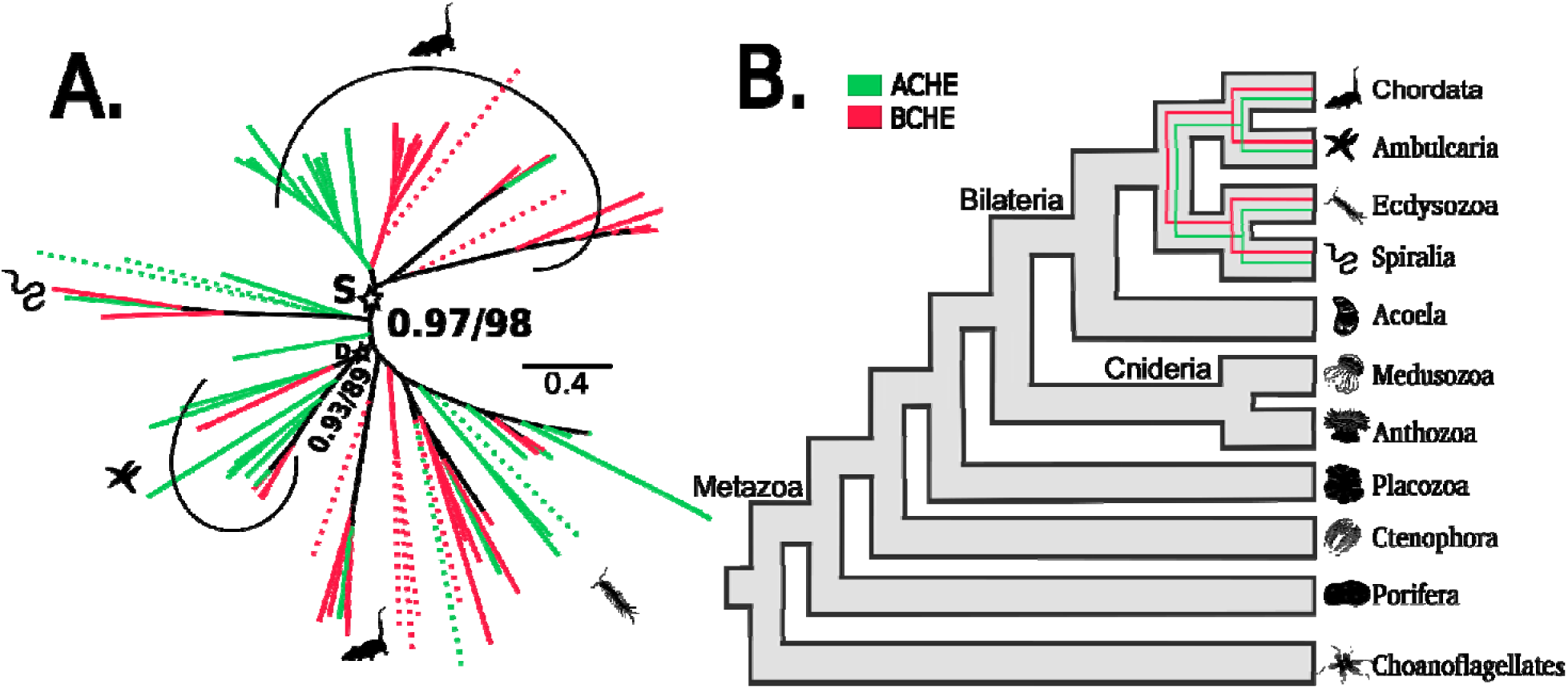
Phylogenetic relationships and gene-tree-species-tree reconciliation of Cholinesterases (ACHE & BCHE). (A) Transfer bootstrap expectation (TBE) phylogenetic trees depicting the evolutionary relationships of the ACHE and BCHE protein families. Nodal support values are shown as transfer bootstrap expectation (TBE; decimal values) and ultrafast bootstrap (UFB; whole-number percentages) for selected nodes. Dashed branches indicate sequences identified as unstable by the t-index or leaf stability index (LSI) analyses. Asterisks (✰) indicate speciation (S) or duplication (D) events, with TBE and UFB support value provided where applicable. (B) Simplified, combined representations of the corresponding gene-tree-species-tree reconciliations inferred using GeneRax. Species silhouettes were obtained from PhyloPic (https://www.phylopic.org/). Complete annotated phylogenetic trees and nodal support values are provided in Online Resource J, and detailed GeneRax reconciliation outputs are provided in Online Resource M(b).

Within vertebrates, human ACHE-associated sequences formed a monophyletic grouping (TBE = 0.96, UFB = 71), whereas BCHE-associated sequences formed a separate, strongly supported grouping (TBE = 1.00, UFB = 97). The node separating these vertebrate-associated groups received comparatively lower support (TBE = 0.89, UFB = 60), suggesting uncertainty in their deeper phylogenetic relationship. The stability assessment identified 13 potentially unstable sequence tips, which were excluded from the subsequent reconciliation, resulting in a dataset containing 99 retained sequence tips.

GeneRax reconciliation under the UndatedDL model identified 61 duplication events and 22 loss events across the sampled cholinesterase-associated dataset (Fig. 3B; Online Resources M(a) and M(b)). The reconciliation mapped the separation of the human-reference-associated ACHE and BCHE groups to a node within the vertebrate portion of the supplied species tree. This placement is consistent with a possible vertebrate-associated duplication scenario, although the comparatively moderate support for the deeper phylogenetic relationship warrants a cautious interpretation. Lineage-associated duplication events were particularly represented in *Anolis carolinensis* (9 events), *Branchiostoma belcheri* (6 events), and *Takifugu rubripes* (5 events). These findings indicate model-inferred diversification within the sampled cholinesterase-associated sequence family, while the precise evolutionary timing and distribution of individual duplication events remain dependent on the gene-tree topology, reconciliation model, and taxonomic sampling.

### RECEPTORS

#### Phylogenetic relationships of muscarinic acetylcholine receptors

Phylogenetic analysis of the recovered human-reference-associated muscarinic acetylcholine receptor sequences comprised 12 terminal sequences from *Homo sapiens, Mus musculus*, and *Monodelphis domestica*, representing the five human receptor subtypes CHRM1–CHRM5. Human and mouse sequences formed strongly supported subtype-associated groupings for CHRM2, CHRM3, and CHRM4 (TBE = 1.00, UFB = 100). CHRM2 and CHRM4 also clustered together with maximal support (TBE = 1.00, UFB = 100), whereas their broader grouping with CHRM3 received comparatively lower support (TBE = 0.82, UFB = 59). For CHRM5, the human-mouse grouping received TBE = 0.90 and UFB = 92, while the broader grouping including the opossum sequence received TBE = 1.00 and UFB = 100. CHRM1 exhibited a less consistent arrangement, with the human and mouse sequences separated from the opossum sequence in the recovered topology.

Despite the support observed for selected subtype-associated groupings, all 12 terminal sequences were identified as potentially unstable under the applied stability criteria. Consequently, these relationships are interpreted as descriptive observations within the recovered mammalian dataset rather than as a resolved reconstruction of muscarinic receptor evolution across Metazoa. No gene-tree/species-tree reconciliation was undertaken for this dataset.

#### Phylogenetic relationships of recovered nicotinic acetylcholine receptor subunits

Using 16 human nicotinic acetylcholine receptor subunit proteins as reference queries, the final phylogenetic dataset comprised 12 terminal sequences representing four human-reference-associated subunit types: CHRNB2, CHRNA6, CHRNA9, and CHRNG. These sequences originated from Homo sapiens, Mus musculus, Monodelphis domestica, and Anolis carolinensis.

CHRNA6 sequences formed a strongly supported subtype-associated grouping across the four sampled species (TBE = 1.00, UFB = 100). Within this group, the grouping comprising opossum, green anole, and mouse sequences received TBE = 0.83 and UFB = 79. CHRNA9 sequences from human, mouse, and opossum also formed a strongly supported grouping (TBE = 1.00, UFB = 100), although the internal human–opossum grouping received comparatively lower support (TBE = 0.54, UFB = 58). The grouping connecting CHRNA6- and CHRNA9-associated sequences received TBE = 0.95 and UFB = 95. Human and mouse CHRNG sequences formed a strongly supported pair (TBE = 1.00, UFB = 100). The CHRNB2 portion of the tree included two human reference entries and one mouse sequence; the two human entries clustered together, while the broader grouping including mouse received UFB = 100.

All 12 terminal sequences were identified as potentially unstable under the applied stability criteria. Thus, although selected subtype-associated relationships were recovered, the limited taxonomic representation and sequence-placement uncertainty restrict their interpretation. These results describe relationships within the recovered human-reference-associated subset and should not be considered a comprehensive reconstruction of nicotinic receptor evolution. No subsequent GeneRax reconciliation was performed for this dataset.

## Discussion

The present study provides a comparative phylogenomic assessment of protein sequences associated with the human cholinergic system across 89 selected eukaryotic proteomes. Using 26 curated human proteins as independent reference queries, we examined the taxonomic representation and phylogenetic relationships of candidate homologues associated with acetylcholine biosynthesis, transport, catabolism, and receptor-mediated signalling. The findings indicate heterogeneous patterns of sequence recovery and phylogenetic diversification among these molecular components. Rather than supporting a uniform evolutionary history for the complete cholinergic system, the results highlight differences in the distributions and evolutionary relationships of its individual protein families.

The CHAT- and SLC18A3-associated datasets showed relatively restricted representation within the sampled proteomes. CHAT-associated candidates were recovered across 24 proteomes, whereas SLC18A3-associated candidates were identified across 19 proteomes, with both datasets represented in multiple bilaterian lineages. These observations are consistent with the representation of the corresponding human-reference-associated protein groups across sampled protostome and deuterostome taxa. However, their apparent non-recovery in several sampled lineages should be interpreted cautiously. Previous molecular characterization has identified CHAT and VAChT homologues in amphioxus and demonstrated their association within the cholinergic gene locus (Candiani *et al*., 2008). Therefore, the failure to recover CHAT from the sampled cephalochordate proteome should be interpreted as non-detection under the present search and filtering conditions rather than definitive evidence of evolutionary loss. Therefore, the failure to recover CHAT from the sampled cephalochordate proteome should be interpreted as non-detection under the present search and filtering conditions rather than definitive evidence of evolutionary loss.

In contrast, the dataset associated with the human SLC5A7 reference sequence exhibited broader taxonomic representation. Candidate sequences belonging to the SLC5 transporter-associated group were recovered from 46 of the 89 sampled proteomes, including representatives of Cnidaria, Ctenophora, and Bilateria. Phylogenetic reconstruction identified a strongly supported vertebrate-associated subgroup containing the human, mouse, and opossum reference-associated sequences. This provides support for the association of the vertebrate subgroup with human SLC5A7. Nevertheless, the broader SLC5-associated dataset exhibited less consistent support across its deeper phylogenetic relationships, including a grouping with UFB = 3 despite TBE = 0.73. Furthermore, the presence of multiple sequence groups within the broader dataset indicates that the recovered sequences should not all be considered specific SLC5A7 orthologues. The inferred duplication and loss events likewise describe diversification within the analysed SLC5-associated dataset rather than confirmed evolutionary events specific to the human CHT1 lineage. The broad distribution of SLC5-family-related sequences therefore highlights the importance of distinguishing transporter-family membership from the evolutionary history of individual human transporter genes.

The cholinesterase analysis revealed a further pattern of phylogenetic differentiation. The recovered sequence dataset included an echinoderm-associated grouping and distinct vertebrate ACHE- and BCHE-associated groups. Within vertebrates, the human ACHE- and BCHE-associated sequences formed separate phylogenetic groupings, although the deeper node separating these groups received comparatively lower support (TBE = 0.89, UFB = 60). Gene-tree/species-tree reconciliation placed the corresponding duplication on a branch within the vertebrate portion of the supplied species tree. This provides a possible vertebrate-associated duplication scenario, consistent with the established differentiation of vertebrate acetylcholinesterase and butyrylcholinesterase lineages (Pezzementi, Nachon and Chatonnet, 2011). However, the moderate support for their deeper phylogenetic relationship means that the precise placement and timing of this separation should be interpreted cautiously. The additional duplication and loss events identified through reconciliation further illustrate the diversification of the sampled cholinesterase-associated sequences, although these model-dependent events should not be regarded as independently confirmed gene-copy changes.

The receptor-associated analyses provided more restricted evidence regarding the relationships among human-reference-associated receptor subtypes. The muscarinic receptor dataset recovered subtype-associated groupings corresponding to the five human receptor reference proteins, with selected relationships receiving strong bootstrap support. Similarly, the nicotinic receptor dataset recovered sequence groupings associated with CHRNA6, CHRNA9, CHRNB2, and CHRNG. Nevertheless, both datasets contained limited taxonomic representation, and all terminal sequences were identified as potentially unstable under the applied stability criteria. Consequently, the recovered relationships provide descriptive information about the sampled sequences but do not independently resolve the evolutionary diversification of muscarinic and nicotinic receptors across Metazoa. This distinction is particularly important because previous comparative studies have documented broader evolutionary diversity of nicotinic receptor subunits, including their presence in cnidarians and substantial lineage-associated expansions in lophotrochozoans (Faltine-Gonzalez and Layden, 2019; Jiao *et al*., 2019). Similarly, previous investigations have examined vertebrate muscarinic receptor diversification in relation to genomic-context changes and early vertebrate genome duplications (Pedersen, Bergqvist and Larhammar, 2018). The restricted recovery observed here should therefore be interpreted within the context of the human-reference-anchored search strategy and subsequent sequence filtering, rather than as evidence of an evolutionarily restricted receptor distribution.

An important consideration in interpreting the present findings is the design of the human-reference-anchored approach. The use of curated human protein sequences provides a consistent framework for examining candidate homologues in relation to defined components of the human cholinergic system. However, this strategy is not equivalent to a comprehensive reconstruction of the complete protein repertoire of every sampled organism. Sequence divergence, lineage-specific gene duplication, incomplete proteome representation, and candidate-filtering criteria can influence the recovery of homologous sequences. In addition, phylogenetic relationships within large and diversified protein families may remain uncertain even when selected terminal groupings receive strong support. The gene-tree/species-tree reconciliation results should consequently be interpreted as model-dependent evolutionary scenarios conditional on the input gene trees, species-tree topology, and sampled sequence representation. In particular, inferred duplication and loss events do not independently establish the exact evolutionary history of individual genomic loci, and non-detection should not be equated with definitive gene loss.

Collectively, the results demonstrate that protein groups corresponding to the human cholinergic system exhibit heterogeneous patterns of taxonomic representation and phylogenetic resolution within the sampled eukaryotic proteomes. The comparatively restricted recovery of CHAT- and SLC18A3-associated candidates, the broader representation of SLC5-family-related sequences, the differentiation of cholinesterase-associated groups, and the limited receptor phylogenies illustrate the importance of examining individual protein families within their respective evolutionary contexts. By applying a common human-reference-anchored analytical framework, this study provides a comparative view of these molecular components and highlights the distinction between broad protein-family relationships and the evolutionary histories of specific human-reference-associated lineages. Further genomic and functional evidence will help refine the evolutionary relationships and biological assignments of these protein groups.

## Conclusion

This study presents a human-reference-anchored comparative phylogenomic assessment of 26 cholinergic reference proteins across 89 selected eukaryotic proteomes. The results reveal heterogeneous patterns of taxonomic representation and phylogenetic relationships among proteins associated with acetylcholine biosynthesis, transport, catabolism, and receptor-mediated signalling. CHAT- and SLC18A3-associated candidates were recovered across multiple sampled bilaterian lineages, whereas SLC5-family-related sequences exhibited broader representation, including sampled non-bilaterian groups. Cholinesterase analyses identified distinct vertebrate ACHE- and BCHE-associated groupings, while the receptor analyses recovered selected subtype-associated relationships within restricted taxonomic datasets. The interpretation of these findings is constrained by the human-reference-anchored search strategy, taxonomic representation, and phylogenetic stability of individual datasets. Overall, this component-wise comparative framework highlights the heterogeneous evolutionary patterns of protein groups corresponding to the human cholinergic system and provides a basis for further investigation of their individual evolutionary relationships.

## Statements and Declarations

## Ethics approval and consent to participate

Not applicable. This study was entirely computational and used only publicly available proteomic and protein sequence data; no human participants, animals, or identifiable private data were involved, so ethics approval and consent to participate were not required.

## Consent for publication

Both authors have read and approved the final manuscript and consent to its submission and publication.

## Authors’ contributions

MK conceptualized the work and AT carried out the research.

## Availability of data, code and materials

All proteomic and protein sequence data used in this study were obtained from publicly available databases, including NCBI and UniProtKB/Swiss-Prot. Supplementary Table S1 provides comprehensive details on the proteomic sequence data, including taxonomic classification, species or proteome counts per taxon, source, accession numbers, and BUSCO completeness scores, as well as the reference sequences used for orthogroup identification. Additional datasets, phylogenetic trees, alignments, and supplementary analyses are available at: 10.6084/m9.figshare.34920324.

